# Scanning electron microscopy of human islet cilia

**DOI:** 10.1101/2023.02.15.528685

**Authors:** Alexander J. Polino, Sanja Sviben, Isabella Melena, David W. Piston, Jing Hughes

**Affiliations:** Department of Cell Biology and Physiology, Washington University School of Medicine, 660 South Euclid Ave, Saint Louis, MO, USA; Washington University Center for Cellular Imaging, Washington University School of Medicine, 660 South Euclid Ave, Saint Louis, MO, USA; Department of Medicine, Washington University School of Medicine, 660 South Euclid Ave, Saint Louis, MO, USA

**Author notes:** Correspondence: Jing Hughes.

## Abstract

Human islet primary cilia are vital glucose-regulating organelles whose structure remains uncharacterized. Scanning electron microscopy (SEM) is a useful technique for studying the surface morphology of membrane projections like primary cilia, but conventional sample preparation does not reveal the sub-membrane axonemal structure which holds key implications for cilia function. To overcome this challenge, we combined SEM with membrane-extraction techniques to examine cilia in native human islets. Our data show well-preserved cilia subdomains which demonstrate both expected and unexpected ultrastructural motifs. Morphometric features were quantified when possible, including axonemal length and diameter, microtubule conformations and chirality. We further describe a novel ciliary ring, a structure that may be a specialization in human islets. Key findings are correlated with fluorescence microscopy and interpreted in the context of cilia function as a cellular sensor and communications locus in pancreatic islets.

## INTRODUCTION

Primary cilia are antenna-like sensory organelles that extend from the centriole and project into the extracellular space, serving important functions in sensory detection and signaling. Since their initial description by transmission electron microscopy (TEM) in the rodent pancreas (*1, 2*), primary cilia have been increasingly examined for their link to human metabolic disease, where altered cilia expression and signaling disrupt pancreatic development and glucose homeostasis (*3–7*). The pancreatic islets of Langerhans are spherical clusters composed of multiple endocrine cell types. Most cells in the islet, as in the rest of the body, project a single primary cilium, its structure most definitively revealed by electron microscopy (*8–12*) in addition to targeted staining by immunohistochemistry. In human islets, primary cilia were first described in beta cells, in 1964 in a case of insulinoma (*13*), whereas validation in non-diseased islet tissues took decades to follow (*14–16*). More recently, islet primary cilia were found to have glucose-regulated motility, suggesting that primary cilia are not simply static sensory organelles (*16, 17*). These findings have fueled growing interest in islet primary cilia in diabetes research, paralleled by an increasing need to understand the basic structure and composition of these cellular projections. What is their shape and size? What is the mechanical basis for their motility? How might their signaling be compartmentalized? Towards answering these questions, we first aimed to determine the physical parameters of islet cilia.

Structurally, primary cilia are divided into several subdomains that are conserved among species and cell types. Lengthwise, cilia are composed of a basal body and an axoneme. The basal body is the mother centriole of the cell, a microtubule-based cylinder that emanates from near the Golgi and remains entirely intracellular. It anchors the primary cilium via a connecting transition zone, a morphologically distinct segment that projects out of the ciliary pocket and controls cargo trafficking to the axoneme. The primary cilia axoneme has a highly conserved conformation of nine outer microtubule doublets with a lack of central microtubules, hence named “9+0,” a pattern consistently observed at the axonemal base and thought to continue through the length of the cilium. This model has required revision, however, due to reports of non-9+0 cilia cross-sections captured by transmission EM as well as 3D tomography of primary cilia in cells of both pancreatic and non-pancreatic origin (*9, 18–22*). The revised view is that the canonical ciliary “9+0” configuration changes soon after leaving the cell body, and that microtubule number and arrangement may be modified along the length of the cilium. We recently reported on the non-9+0 arrangement of human islet cilia, but only in sporadic thin sections of resin-embedded tissue prepared for TEM, while how the axoneme structure evolves from base to tip is unknown (*16*). Many other morphometric features of human islet primary cilia also remain undefined, including axonemal length and diameter, geometry of microtubule filaments, whether they assemble higher-order conformations, and the presence of any accessory components contributing to the maintenance of axonemal integrity, dynamics, and sensory signaling function. Thus, elucidating cilia structure and especially visualizing the entire axonemal cytoskeleton is a priority in islet cilia research.

Advances in super-resolution and electron microscopy combined with increasing availability of human donor specimens makes it now possible to perform ultrastructural studies on primary human islets. Large open-access databases such as the nPOD nanotomy atlas have curated scanning transmission EM images of human islets (*23*). While these images are useful for demonstrating disease-related morphological alterations across cellular compartments, cilia and centrosomes were not examined in these studies owing to their rarity and low likelihood of identification. To enable ultrastructural characterization of islet primary cilia, we used surface topography imaging by scanning electron microscopy (SEM) to directly visualize the ciliary axoneme in human pancreatic islets. This approach allows cilia identification based on their extracellular localization and distinct morphology and preserves full-length cilia structure *in situ*. We examined cilia with or without demembranation to compare surface versus cytoskeletal details of primary cilia, both contextually within intact islet tissues (**Figure 1**). Ciliary structures post-demembranation were preserved with clarity, allowing quantification of microtubule size, number, chirality, and observation of structural motifs not previously described in other mammalian cilia.

**Figure 1.**
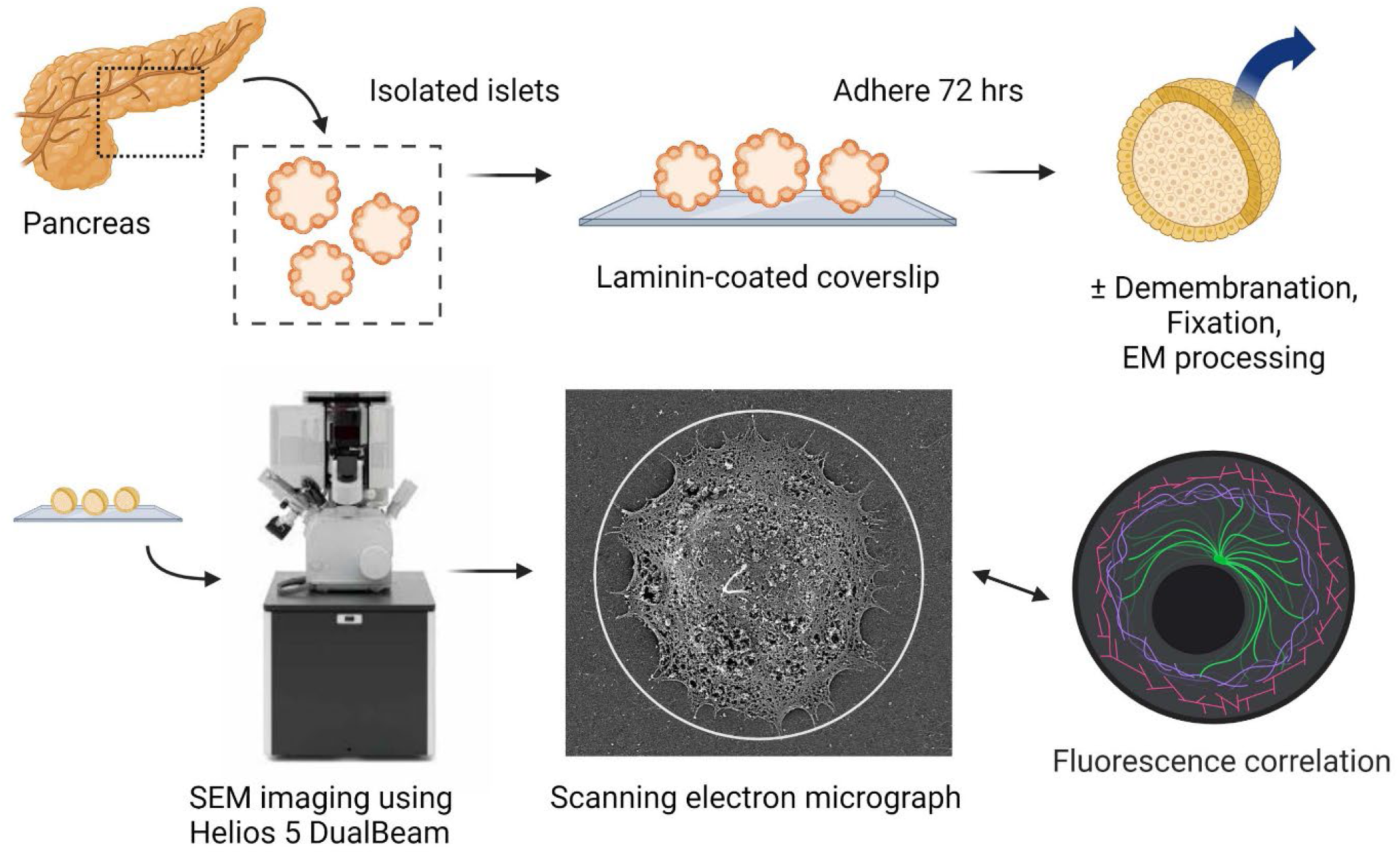
Scanning EM workflow for visualizing primary cilia on intact human islets.

## RESULTS

We examined individual cilia on 15 islets from 3 human donors (**Table 1**). The islets in our study were isolated from human donor pancreata and ranged in size from 100 to 300 μm in diameter, consistent with dimensions reported in literature. Intact islets exhibited round shape and well-preserved surface contour, with the contrasting textures among adjacent cells exhibiting a soccer ball-like appearance (**Figure 2a**). Primary cilia are prominent structures on the islet surface, measuring 3 to 6 μm in length, are flexible and each displaying varying curvature. Other membrane protuberances such as microvilli and filopodia are also observed among islet cells. As expected, the vast majority of human islet cells bear a single cilium each, a feature that we and others have previously demonstrated by immunofluorescence microscopy (*14, 16, 24, 25*)

**Table 1.** Human islet donor characteristics.

| Donor | Gender | Age (y) | BMI (kg/m<sup>2</sup>) | Diabetes |
| --- | --- | --- | --- | --- |
| 1 | M | 54 | 28.4 | No |
| 2 | M | 32 | 26.9 | No |
| 3 | M | 16 | 29.5 | No |

**Figure 2.**
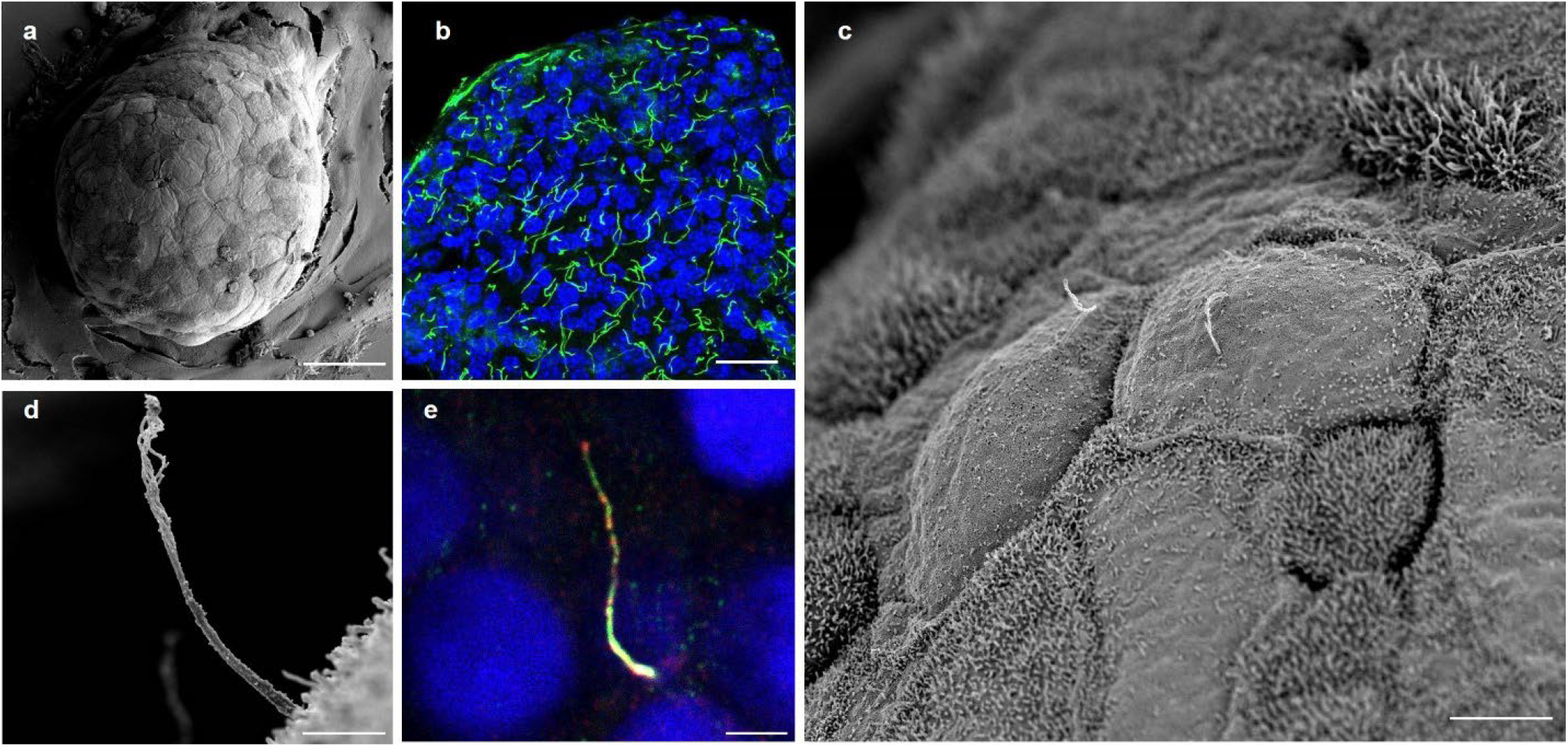
Multi-scale imaging of human islet primary cilia. (**a**) Intact human islets are immobilized on glass coverslips and membrane-extracted to reveal ultrastructural surface details. Shown is an average-sized islet with approximately 100 cells on the exposed surface. Scale, 50 μm. (**b**) Confocal image of a human islet showing greater density of primary cilia seen through a projected z-stack. Acetylated-α-tubulin, cilia (green), DAPI, nuclei (blue). Scale, 20 μm. (**c**) Higher-resolution view of an islet region showing two centrally-located cells projecting primary cilia. Surface texture differences from the presence or absence of microvilli are visible among adjacent cells. Scale, 10 μm. (**d**) A full-length cilium is captured in profile on the islet surface, showing evolving morphology from proximal to distal segments. Scale, 2 μm. (**e**) A human islet cilium stained by ciliary markers ARL13b (red) and axonemal dynein DNAI1 (green), showing comparable cilia length, width, and curvature as those in SEM images. Scale, 2 μm.

In the present study, primary cilia were imaged with or without de-membranation, the latter revealing a clear view of the axonemal structure from base to tip. In total 34 full-length cilia including 28 membrane-extracted and 6 non-extracted samples were selected for morphometric analysis. Multi-scale SEM was used to visualize structural details of each cilium and its sub-domains with progressively higher resolution.

### Ciliary length, diameter, and volume estimations

Most cilia on the islet periphery project away from the cell, while some project horizontally along the membrane (**Figure 3**). We sought to determine the linear dimensions of islet primary cilia, keeping in mind the caveat that measurements of 3D object length in 2D images are by nature underestimations. By virtue of the object tilt, planar-extrapolated lengths can only be equal to or shorter than actual object length. To achieve the best estimation of ciliary length, we selected mostly cilia that were captured in profile perpendicular to the imaging detector. Average axonemal length was 4.71 ± 1.5 μm among 28 extracted cilia and 4.80 ± 1.5 μm among 6 unextracted cilia, with similar distributions (**Figure 3c**). As human islet cilia are wider at the base than tip, we measured ciliary diameter both proximally and distally, obtaining an average of 225 ± 91 nm at the base and 138 ± 41 nm at the tip for extracted cilia. These dimensions were slightly greater for unextracted cilia, 260 ± 103 nm at the base and 142 ± 60 nm at the tip. Overall length and diameter measurements were comparable to those determined by cryo-electron tomography (cryo-ET) in cultured kidney cell cilia and those reported for other mammalian cilia (*19–21*).

**Figure 3.**
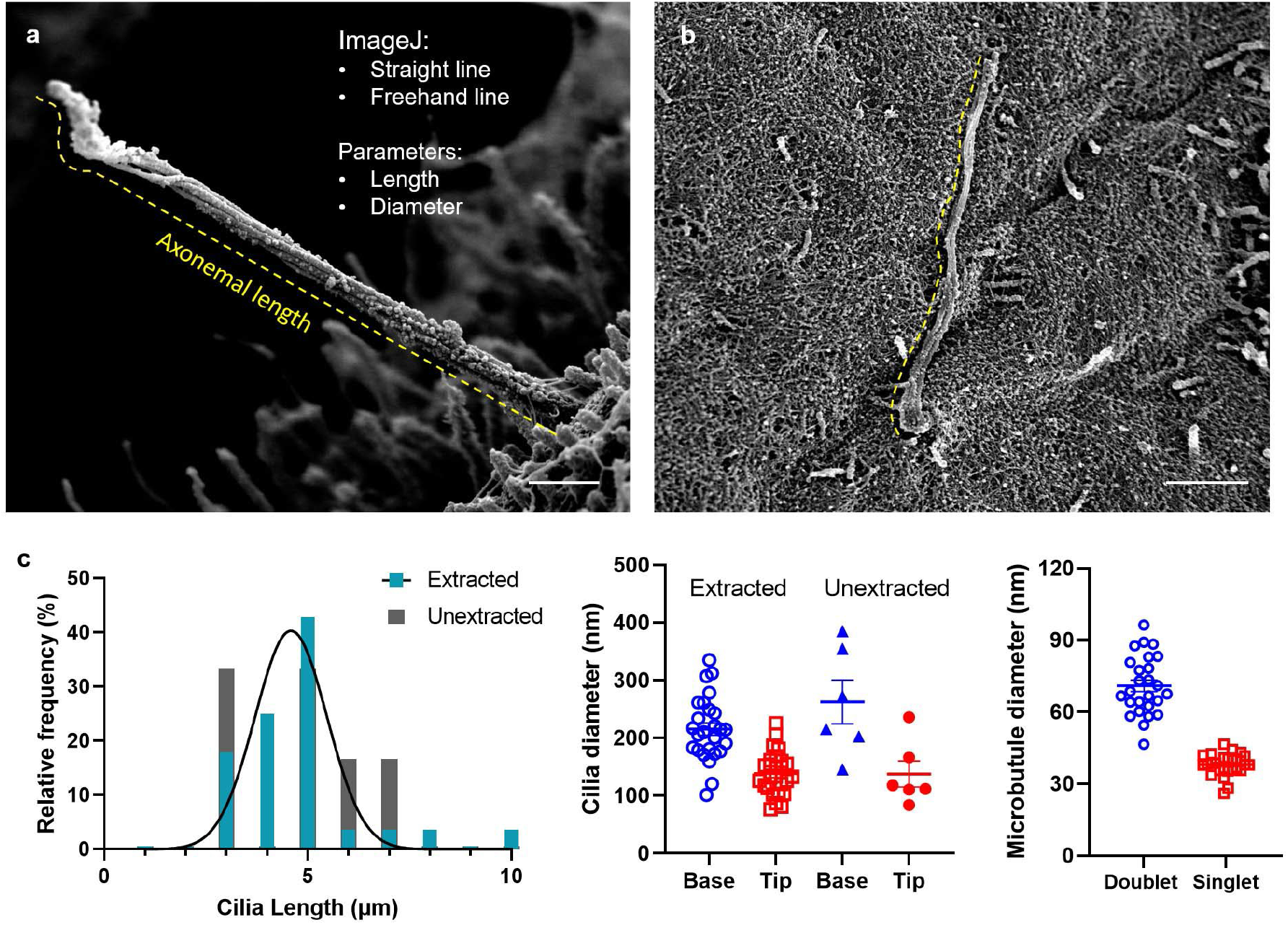
Cilia length measurements at the islet surface. Full-length cilia with intact tips are used for axonemal length determinations. (**a**) A cilium projecting away from the islet surface with a relatively straight axoneme, whose length can be measured using a combination of the straight line and freehand line tracing tool in ImageJ, shown in yellow dash. Cross-axonemal diameters can be measured using a straight line tool at the cilia base and tip. Scale, 500 nm. (**b**) A curvy cilium along the islet surface, whose length can be traced using the freehand line tool. Scale, 2 μm. (**c**) Morphometric analysis of human islet cilia showing distribution of axonemal length among all 34 primary cilia (n= 28 extracted, 6 unextracted), base and tip diameters, and microtubule doublet versus singlet diameter in extracted samples. Bars represent mean ± SD.

Using measurements from unextracted cilia, which had intact membranes, we estimated the human islet primary cilium volume and surface area based on the shape of either a truncated circular cone or a right cylinder (**Figure 4**). As a cone with a base and tip diameter of 0.26 and 0.14 μm (radius 0.13 and 0.07 μm), respectively, the cilium volume is estimated to be 0.16 μm^3^, with a surface area of 3.10 μm^2^. As a cylinder, with an average axonemal diameter of 0.2 μm (radius 0.1 μm), the cilium has a volume of 0.15 μm^3^ and surface area of 3.08 μm^2^. Thus the two model methods were in good agreement with each other.

**Figure 4.**
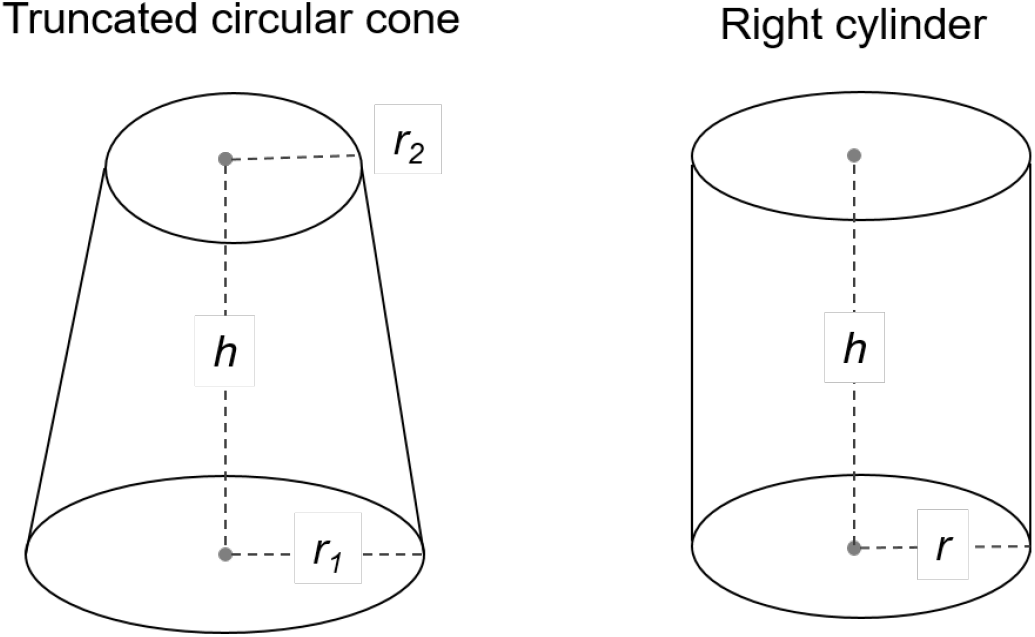
Geometric models of primary cilia. The shape of human islet primary cilia can be modeled as either a truncated circular cone or a right cylinder. These simplified representations allow calculations of volume and surface area using empirically determined radius and height measurements.

We next measured the average islet cell diameter in our dataset, which was 12-14 μm (radius 6-7 μm), and determined the cytoplasmic volume to be ~1,000 μm^3^. This allowed us to calculate the ratio of cytoplasmic volume to ciliary volume for human islet cells as approximately 6,000-7,000. The ratio of plasma membrane surface area to ciliary membrane area is about 150-200. These ratios were previously estimated to be 5,000 and 500, respectively, for a typical 0.3 × 5 μm primary cilium (*26*). The discrepancy between our and previous estimates, as well as the fact that our ratios show greater contrast in volume than surface area, results from the fact that human islet cilia have smaller diameter than that generally assumed (0.2 μm versus 0.3 μm). In other words, human islet cilia are more slender than what may be typical for other mammalian primary cilia. Nonetheless, the values we obtained for human islet cilia are within a 2-fold range of prior estimates using non-islet models (*26, 27*), indicating that the physical dimensions of primary cilia are largely conserved across cells and organisms.

The actual shape of the human islet primary cilia, of course, is not a uniform cone or cylinder. Our SEM imaging revealed remarkable cilia geometries and structural elements, which we now describe from base to tip. Special attention is given to axonemal domains with known functional importance, as well as previously uncharacterized structural motifs.

### The ciliary pocket and transition zone are well-visualized by SEM

The ciliary pocket is an invagination of the plasma membrane from which the primary cilium emanates. It is a specialized domain for ciliary protein trafficking, endocytosis, and actin cable docking, among other functions (*28, 29*). Ciliary pockets had been observed in mouse islets more than a half century ago by TEM, in beta cells exhibiting deeply rooted cilia where the ciliary pocket was interpreted as a site of basal body docking to the plasma membrane, with a presumed role in primary ciliogenesis (*1*). In the present study, the ciliary pocket of human islets is seen to form a circular pit surrounding the axoneme in some cells, while in others the cilium erupts from the surface without an appreciable ciliary pocket, and there are other cells still that have such deeply rooted cilia within recesses of the cell that a ciliary pocket is not visible (**Figure 5**). From these observations we conclude that the ciliary pocket adopts specialized configurations among different human islet cells, a theme that is consistent with the differential membrane organization at the cilia base that has been described in other systems (*28–31*).

**Figure 5.**
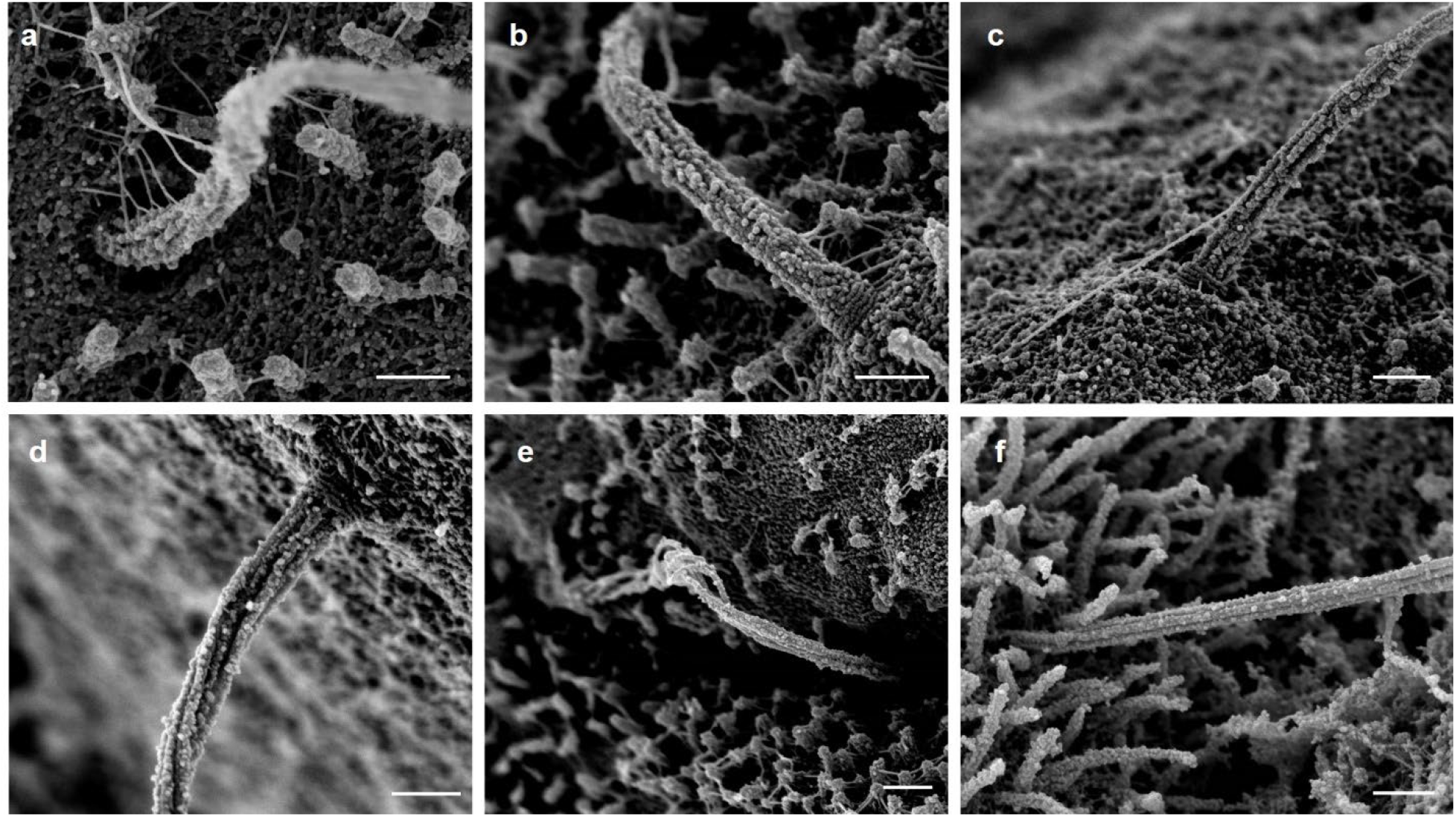
Heterogeneous morphology at the ciliary base. The proximal region of human islet cilia differs among cells, showing variable depths of ciliary pocket and basal attachments (**a-d**) including Y-shaped linkers between the cilium base and nearby surface structures. In other cilia, the base is buried within membrane recesses and microvilli and therefore not visualized (**e-f**). The ciliary necklace, when exposed, can be visualized in exquisite detail (**b-d**), revealing discrete rows of particles in the transition zone that are perpendicular to the axonemal microtubules. Scale, 300 nm.

Higher up, the transition zone is the proximal-most region of the axoneme and is captured in striking clarity in our study. There is a section of 5-6 parallel rows of particles that encircle the cilia base orthogonal to the direction of axonemal microtubules (**Figure 5b-d**), corresponding to the “ciliary necklace” that has been described at the neck of the cilium in many species, where it supports the microtubule transition from triplets to doublets and acts as a membrane diffusion barrier (*32–36*). The necklace in human islet cilia measures approximately 230 nm in diameter and 140 nm in height in the visible portion above the cell. Individual bead-like particles comprising the necklace measure approximately 12.5 nm in diameter each, similar to the dimensions of the intramembrane particles (IMPs) previously described for kinocilia in the respiratory epithelium (*37*). The distance between the necklace rows, or strands, is about 18-22 nm, and the number of extracellular strands ranges between 3 and 6 per cilium. Whereas the ciliary necklace had best been previously demonstrated by freeze-etching EM (*32, 37*), a technique that relies on a carbon cast replica of the structures in study, our imaging approach visualizes the structure itself, allowing direct morphometric measurements of its components.

Distal to the ciliary necklace, we see cylindrical strands of microtubule doublets forming the ciliary axoneme. Small knob-like densities with diameters of ~20 nm are observed decorating the microtubules and are similar in size and shape as the beads that constitute the ciliary necklace (**Figure 5**). These particles are also present on microvilli and cortical cytoskeleton, in a dense albeit irregular way, thus appear non-specific to cilia and may represent membrane remnants or debris post-extraction and post-fixation. Occasional larger appendages are seen to associate with the axoneme surface (**Figure 6a**), but we did not observe regular repeats of polymer complexes resembling intraflagellar transport (IFT) trains (*19, 42*) which may not have survived demembranation.

**Figure 6.**
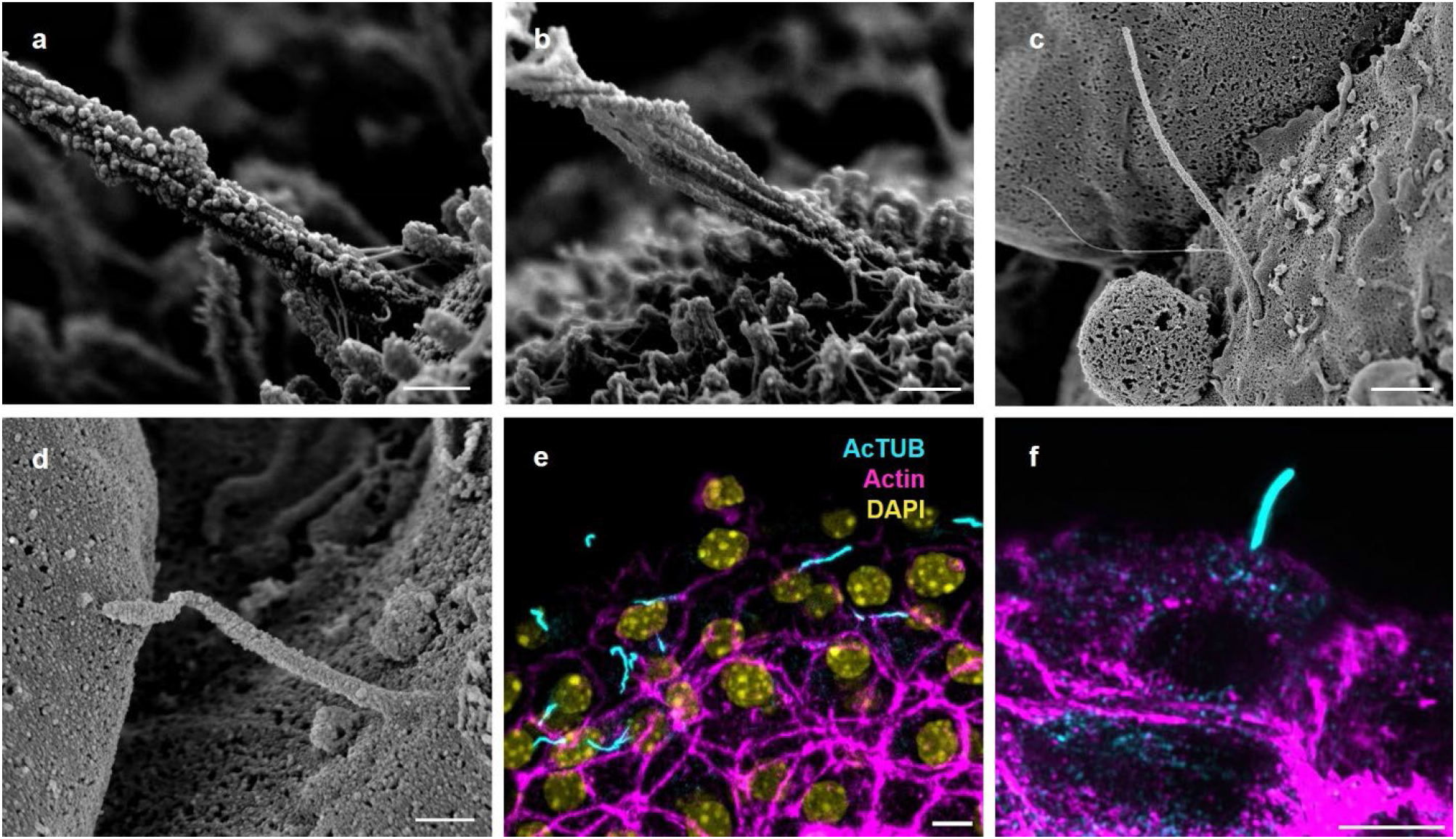
Actin-like microfilaments attach at the cilium base. Thin filaments connect the base of the cilium to microvilli and the cell surface; branching patterns are seen at the axonemal and microvilli attachment points (**a-b**); scale 300 nm. These microfilament tethers are not visible in unextracted islet samples that have preserved plasma and ciliary membranes (**c-d**); scale, (**c**) 1 μm, (**d**) 400 nm. (**e-f**) Actin filaments are visualized by phalloidin staining (magenta) at the base of primary cilia (cyan); nuclei (yellow), scale 5 μm, structured illumination microscopy.

### Cilia are tethered by microfilaments at the base

We observe physical interactions between the ciliary axoneme and ciliary pocket as well as with the cell surface. The base of the primary cilium is connected by thin filaments to the surrounding microvilli and cytoskeleton surface (**Figure 5a-c, Figure 6a-b**). These microfilaments have a diameter of 7-10 nm and a branching form, consistent with actin fibers, which are known to associate with primary cilia – particularly at the base (*28, 38–40*). Location and morphology-wise, they are distinct from the Y-links in the transition zone of cilia and flagella (*21, 32, 41–43*) which are symmetrically present on all microtubules and serve to tether the microtubules to the ciliary membrane. In contrast, these external filamentous strands observed in our study are located above the transition zone, attach to distant targets including neighboring microvilli and cell surface, and are asymmetrically arranged around the axoneme, predominating on one side of the cilium (**Figure 5a-b**). Similar filaments are also present on microvilli, where they are more symmetrically distributed and create an aster-like network. Neither cilia-nor microvilli-associated filaments are visible in non-demembranated samples (**Figure 6c-d**), suggesting that these structures are normally sheathed by the ciliary and plasma membranes. Fluorescence imaging of mouse islets using structured illumination microscopy (SIM) provides supporting evidence that the base of primary cilia is embedded in an F-actin network (**Figure 6e-f**).

### Microtubule doublet-to-singlet transition

In all observed cilia, individual microtubule filaments start as doublets at the cilia base and transition to singlets at the one-third to half-way mark along the axoneme, between 1-2 μm from the base, at which point the cilium takes on a noticeable change in diameter and width (**Figure 7**). There is a clear demarcation where this texture change takes place, beyond which point the microtubules become more loosely associated, reflecting a change in filament organization and intra-filament crosslinking. Such microtubule doublet-to-singlet transitions have been reported in cultured kidney cell cilia (*19, 21*) and can also be observed in open-access volumetric imaging of rodent and human pancreatic islets (*20, 22, 23*). Direct measurements from our dataset showed that human islet cilia microtubule singlets possess an average diameter of 37 ± 5 nm, almost exactly half that measured of microtubule doublets (72 ± 13 nm). These numbers are consistent with those previously reported in kidney primary cilia (*19, 21*). Small globular densities are seen coat the microtubule doublets in the proximal axoneme, whereas they are generally sparse or absent on distal portions of microtubules (**Figures 7-8**).

**Figure 7.**
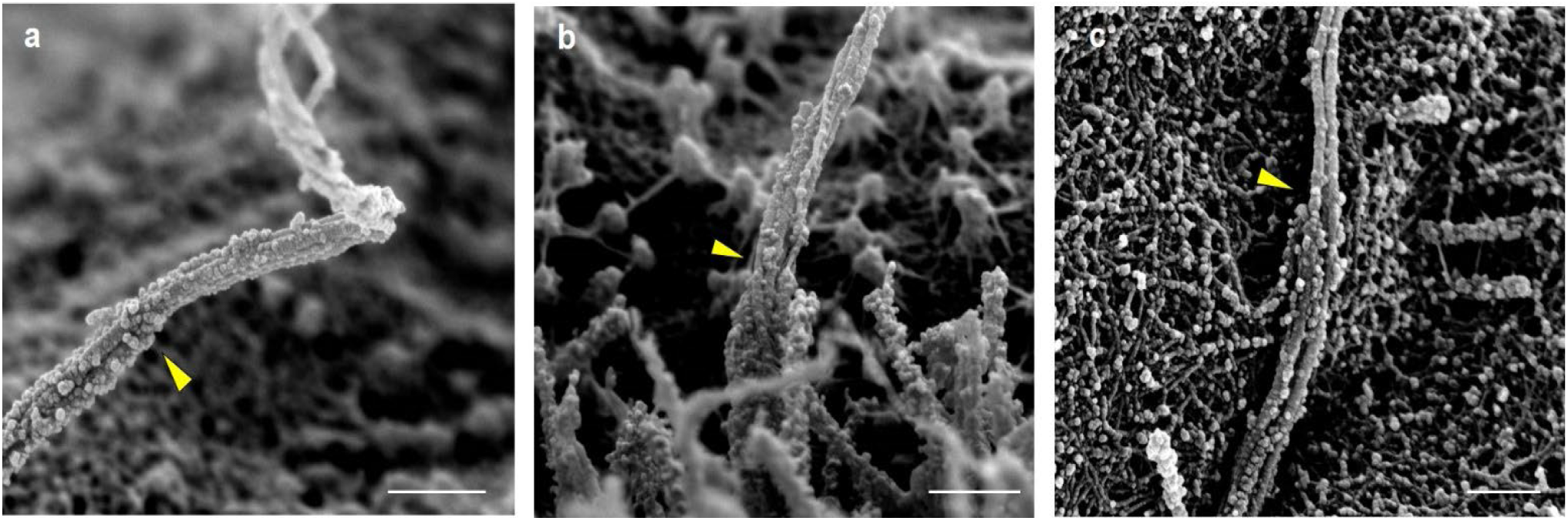
Microtubule doublet-to-singlet transitions are visible on the axoneme. Representative images showing the demarcation along the axoneme where microtubule doublets become singlets, resulting in a reduction of microfilament and axoneme diameter. Transition points marked by yellow arrowheads. Scale, 300 nm.

**Figure 8.**
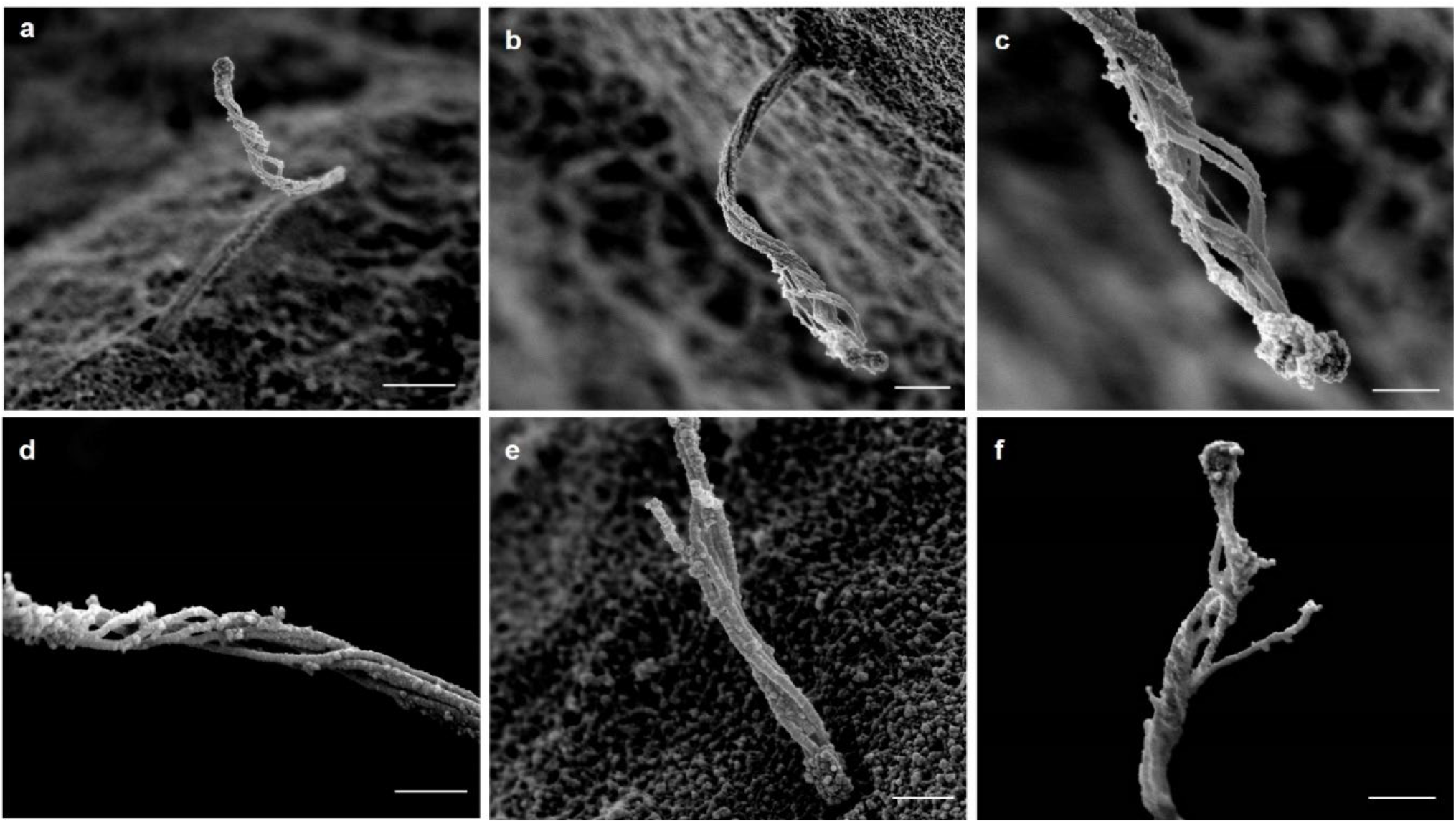
Microtubule configurations in the distal axoneme. A survey of different axonemal configurations in the distal region. Microtubule both rotate and separate in the distal third of the axoneme, creating a helical cage. Occasional actin-like thin filaments are seen among the microtubules (**c**). Splaying of individual microtubule filaments may be secondary to specimen damage (**e-f**). Scale, **a-b**, 500 nm; **c-f**, 300 nm.

### Cilia handedness and secondary structure

As microtubule filaments evolve from doublets to singlets, they undergo conformational changes with respect to each other. Whereas they begin as a tight straight bundle in the ciliary baso-proximal region, the microtubules begin to separate distally and twist about a central axis before coalescing at the tip (**Figure 8**). There is a periodicity to the microtubule rotations, measuring 900-1000 nm in arc length with a pitch of 500-1000 nm between turns. Within the loose microtubule bundle, actin-like thin filaments with a diameter of ~8 nm are occasionally observed stretching along the long microtubule axis (**Figure 8c**). Occasional microtubules break or splay apart (**Figure 8e-f**), likely as a consequence of physical disruptions from extraction and fixation. Given its helical nature, the axoneme is intrinsically chiral. Human islet cilia are predominantly left-handed (>80%), while a minority have a mild right-handedness (~10%), and another small number of cilia have no discernable rotation (**Figure 9**). This is in contrast to the predominant right-handed spins reported for kidney cell cilia (*19, 21*). To be sure of our observations, we confirmed using fiducial markers that our SEM images are not mirror images. The presence or absence of taxol also did not alter the rotational nature and direction, suggesting that these higher-order microtubule conformations are fairly stable.

**Figure 9.**
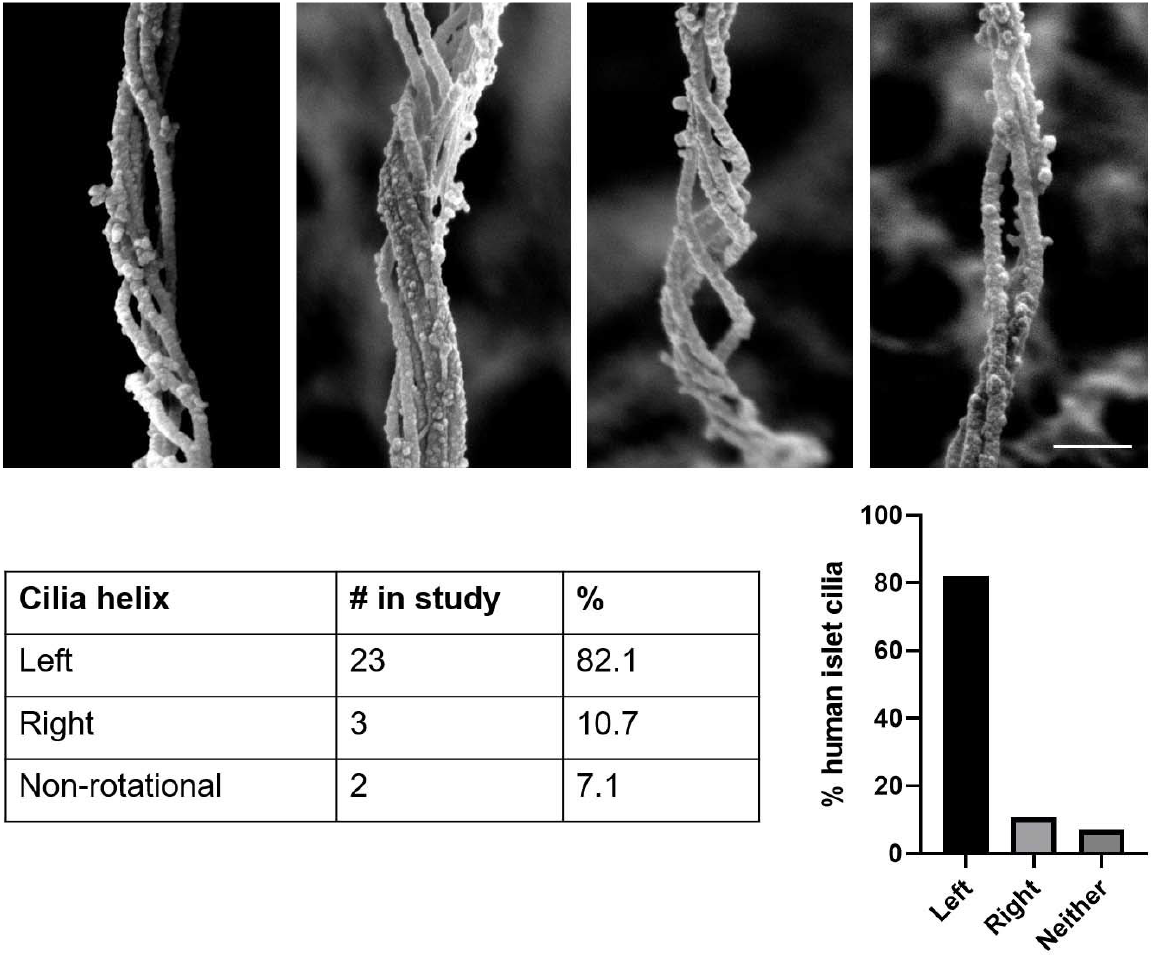
Cilia handedness in human islets. Primary cilia in human islets exhibit chirality in axonemal microtubule arrangements, examples shown from mid-distal axonemal regions where rotations are most pronounced. Quantitation among 28 extracted cilia shows that the majority of islet cilia have left-handed turns. Scale, 200 nm.

### A ciliary ring is occasionally observed in the proximal axoneme

We observe a rare but potentially important feature which is that a ring is present in the proximal axoneme, at about 1-1.5 μm from the base. It wraps around the axonemal bundle as a continuous strand with similar texture and diameter as a microtubule singlet or doublet (30 or 60 nm) (**Figure 10**). We name this structure the “ciliary ring” to differentiate it from the ciliary necklace and ciliary bracelet, which are separate structures (*32, 44, 45*). Rings are observed on 4 out of 34 cilia (12%) from multiple donors, thus not an isolated phenomenon. All rings occurred on cilia that were surrounded by abundant microvilli and therefore likely on beta cells. One cilium was observed to bear two rings that subdivide the axoneme into equidistant thirds (**Figure 10c-e**). Distal to each ring, the axoneme exhibits greater separation and twist, appearing as if the ring is acting as a band to keep the microtubule bundle together. Rings are not observed on non-extracted cilia, indicating that the structure occurs sub-membrane.

**Figure 10.**
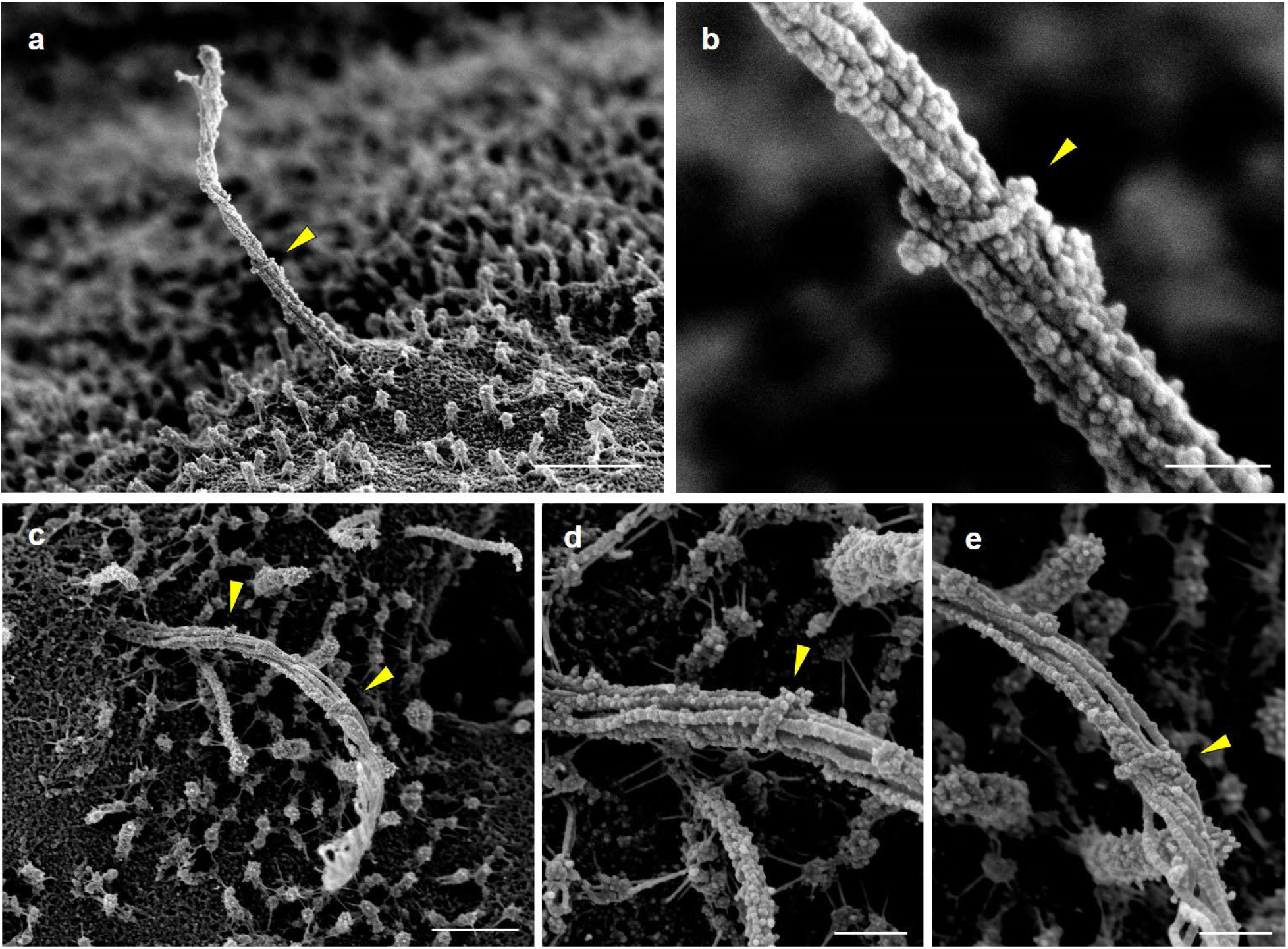
A ciliary ring encircles the axoneme. Ring-like structures found on human beta cell cilia, shown here from different vantage points and marked by arrowheads. Note the contrasting cilia morphology proximal vs. distal to the ring, where the ring location marks where the axonemal bundle begins to unwind and twist. **(b, d, e)** are high-magnification views of (**a, c)**. Scale, (**a, c)** 1 μm; (**b, d, e**) 200 nm.

### Ciliary tip as a special domain

The distal ciliary domain, or tip segment, is captured intact in demembranated islet cells, showing structural organization that differs from the rest of the axoneme. The microtubules decrease in number towards the distal cilium before terminating into dense cap structure (**Figure 11a-c**). There is diversity in the number of microtubules that make it to the tip, typically between 3 and 5, but up to 7 have been counted. A greater diversity in surface morphology can be seen in non-extracted samples in which the cilium retains its membrane covering, where the enclosed tip shape ranges from pointed to rounded to bulbous (**Figure 11d-f**). The highly dilated tip seen in **Figure 11f** resembles those previously described in normal kidney collecting duct cilia (*46*) as well as cilia bearing mutations affecting retrograde transport that cause a buildup of cargo and vesicles in the tip region (*47–49*). Meanwhile, our specimens came from healthy donors with no known ciliopathies, suggesting that these morphological differences may represent a normal spectrum of pancreatic islet cilia in various stages of growth and remodeling. Overall, the unique morphology of the cilia tip observed in our study is consistent with its role as a dedicated sensory and signaling domain, where the diversity of tip structure may be related to specific ciliary roles in different islet cells.

**Figure 11.**
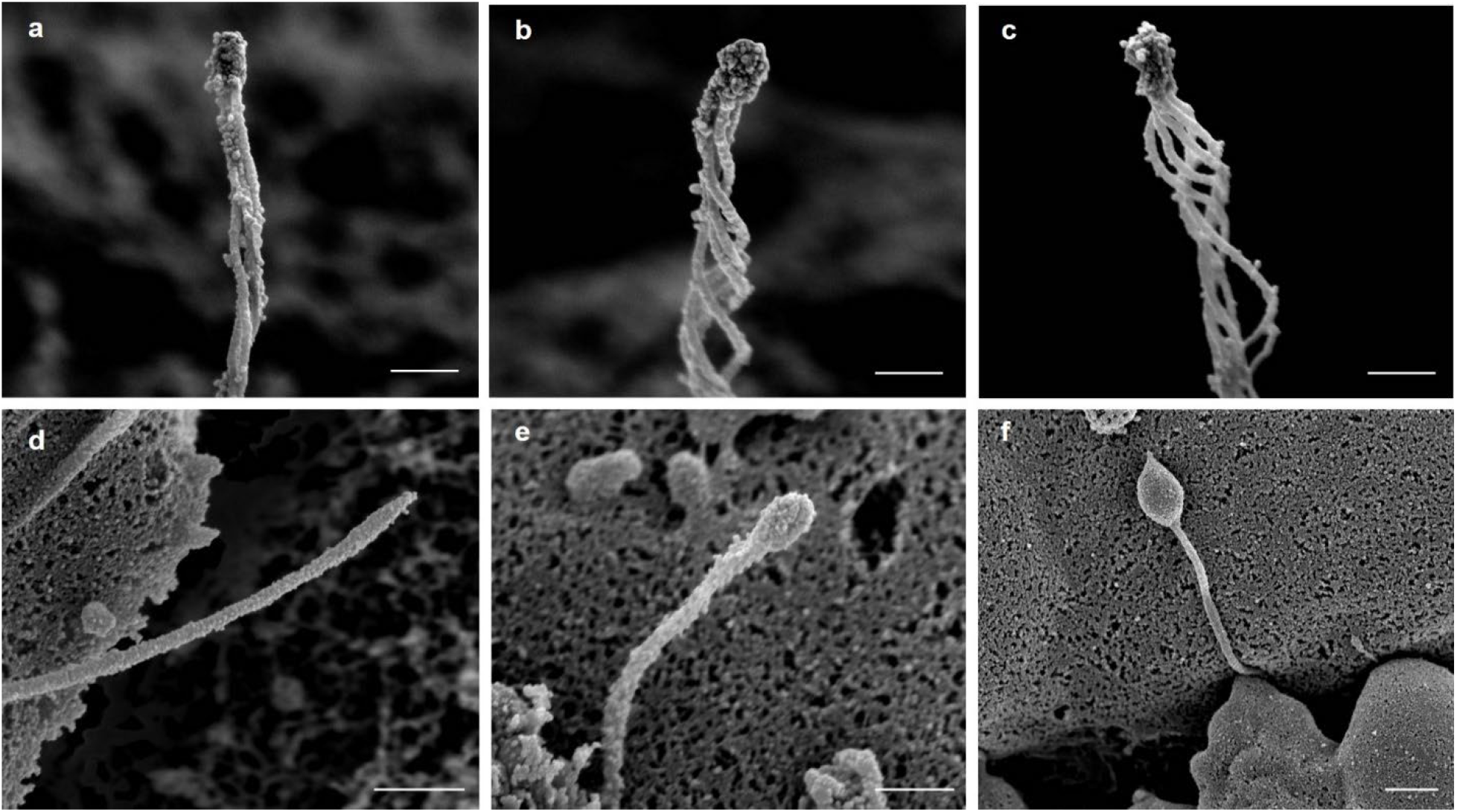
Diversity of cilia tip morphology in human islet cells. (**a-c**) Demembranation reveals a conserved central microtubule organization where the microtubule filaments coalesce and terminate at a dense tip complex. The number of microtubules present at the tip is usually between 3 and 5. (**d-f**) Unextracted samples with intact ciliary membrane show greater heterogeneity including ciliary bulbs and bulges. Scale, **a-c**, 300 nm; **d-f**, 400 nm.

### Multi-ciliated islet cells

Occasional compound cilia are observed by SEM where a single cell may project two or even three cilia from the same base (**Figure 12**). These are rare, estimated <5% frequency among human islet cells. We observed 3 cases of double cilia and 1 case of triple cilia. One clear example each is shown below from unextracted cells that had intact cilia membrane. In both cases, the double and triple cilia projected axonemes that were similar in length and diameter and exhibited varying degrees of physical interactions amongst the axonemes. These project as independent axonemes, analogous to the human dermatologic condition of *pili multigemini* or “tufting” where multiple hair fibers emerge from the same follicle (*50*). As with differences observed in cilia tip morphology, these multi-cilia may represent either normal heterogeneity or aberrant variations among islet primary cilia, as the mechanisms governing centrosome duplication and ciliogenesis are yet unclear in human islets.

**Figure 12.**
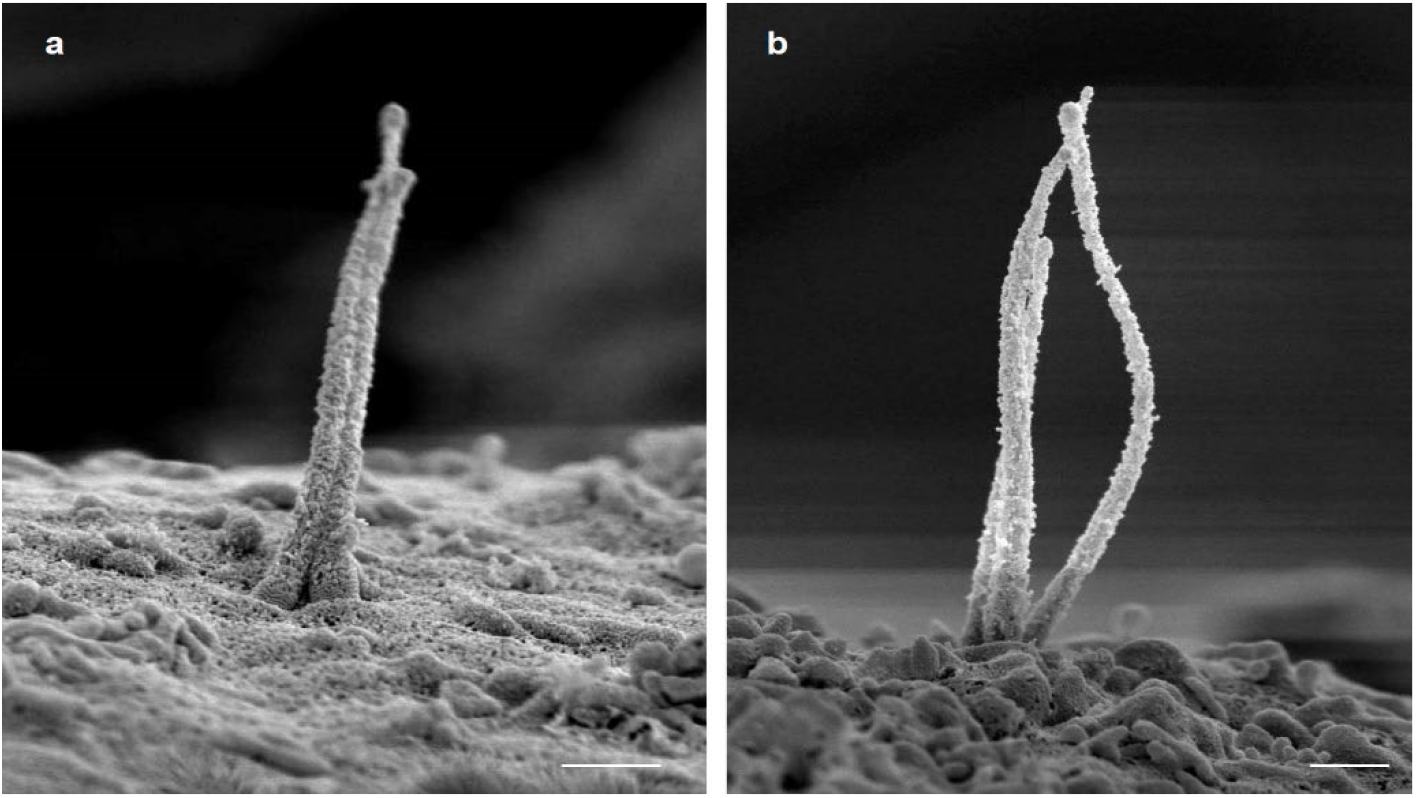
Multi-cilia in human islet cells. (**a**) Double cilia emanating from a single enlarged pocket with two parallel axonemes similar in length and diameter, both possessing defined tips. (**b**) Three axonemes sharing a connected base, physically uncoupled but interactive in their tip regions. Scales, 500 nm. Minor background distortion is seen in both images from charge build-up during scanning.

## DISCUSSION

Our data provide the highest-resolution view to-date of human primary cilia in any tissue and organ system. Recent advances in volumetric electron microscopy including focused ion beam scanning electron microscopy (FIB-SEM) and electron tomography (ET) has allowed 3D examinations of mammalian primary cilia including in rodent islets (*15, 20, 22*), however no human islet data is available from these approaches. Our study fills this important gap, combining 3D approaches to directly visualize cilia *in situ* and multi-scale imaging to resolve structural details, allowing characterization of the full ciliary axoneme from multiple human donor islets. Some key limitations and implications of our work are discussed below.

An important question that we could not formally address is the heterogeneity of cilia morphology among islet alpha/beta/delta cells. The identification of islet cell types, while a simple task by light microscopy and TEM, are much trickier on SEM as there are not many identifying features unequivocally established for endocrine cell groups. We attempted to classify cells by surface morphology, relying on ultrastructural features reported by previous studies of human islets (*23, 51, 52*). For example, beta cells are known to express abundant microvilli (*53, 54*) which corresponded to a characteristic velvety appearance on their surface (**Figure 2, 10**). Alpha cells, in contrast, have smoother surfaces that are largely devoid of microvilli (**Figure 2c, 5d**). Delta cells in human islets tend to be elongated and form long filopodial extensions (*55*), but whether they possess microvilli and whether these correspond to the much longer villi (**Figure 5f, 7b**) than those observed on beta cells remains a question. Cellular abundance and distribution offer helpful clues, as the surface of human islets contains intermingling beta, alpha, and delta cells in roughly 5:3:1 ratio (*51*). Among the 34 cilia that were sampled in entirety in our dataset, we identified 20 beta, 8 alpha, and 3 delta cells based on the postulation that beta cells express abundant short microvilli, alpha cells have few or no microvilli, and delta cells are the rare cells with long villi. This was a limitation in our dataset that can be addressed by future studies, where true cell identity might be ascertained by immuno-SEM or correlative light and electron microscopy (CLEM) using hormonal or cell surface protein markers. Elemental analysis using energy dispersive X-ray spectroscopy (EDS/EDX) is another possibility, which could be used to identify islet cell type by relative abundance of phosphorus, sulfur, and nitrogen, corresponding to alpha, beta, and delta cell secretory granules (*23, 52*).

The arrangement of actin-like connections at the ciliary base offer clues to the role of actin in cilia dynamics. The ciliary pocket has been described to be a docking site for actin cables (*28, 56, 57*), where interactions between the actin cortex and the cilium may serve to either stabilize cilia or actually generate forces to influence cilia movement (*39, 58, 59*). The asymmetry and, sometimes one-sidedness, of these fibers observed in our SEM data (**Figure 5a-c**) suggest that they may act as ropes to pull the cilium via extraciliary forces, similar to a model proposed by Lippincott-Schwartz and colleagues (*39*). Such forces may contribute to primary cilia motility in addition to microtubule-dynein activities (*16, 39*). If, on the other hand, these actin-like filaments were for stability purposes, one would expect them to surround the base in more uniform or symmetric fashion to prevent axonemal fluctuations. Other potential functions of actin and associated myosin at the cilia base may include regulation of ciliogenesis, establishment of cilia orientation, and regulation of endocytic uptake, as demonstrated in several experimental systems (*58, 60–64*). The observation of actin-like filaments along the central axonemal axis (**Figure 8c**) suggests that ciliary actin could be a constitutive component that either provides mechanical stability to the axoneme or partitions the cilia into functional microdomains, as it has been proposed for non-islet cilia (*19, 38*), thus it would be informative to test these models experimentally in islet cells.

The helical nature and chirality of axonemal microtubules was an unexpected feature that may have functional significance. Our finding that islet cilia are mainly L-handed helices contrasts with earlier reports in kidney cell cilia which produce mild R-handed spirals (*19, 21*). In our unpublished TEM observations, centriolar triplets in human islets exhibit an anticlockwise spin, consistent with its consensus orientation in other tissues (*65–67*). If such a rotation were continued along microtubule doublets and singlets, this would indeed translate to a L-handed axoneme. While the functional difference of L-versus R-handed turns is yet unclear, we could at least conclude that cilia handedness *can* differ among cells, and speculatively, we may propose that it may differ among the same cell type but under different growing conditions, a phenomenon well-described in the *Arabidopsis* plant (*68*). The more fundamental question is, what is the function of coiling? In contrast to individual tubulin protofilaments in which curved tips are thought to facilitate growth (*69*), collective coiling by the microtubule bundle likely contributes to overall axonemal stability and dynamics. With the microtubules fixed at both base and tip, the ciliary axoneme can be considered a topologically closed system, where changes in its helical conformation would allow elastic deformation without breaking strands of the microtubule bundle. In other words, the helical unwinding would be a quick way to increase ciliary length without polymerizing new subunits. Additionally, having loose microtubule helices would produce greater cilia surface area and luminal volume, which may alter the capacity for ciliary signaling, cargo trafficking, and exocytic vesicle release. However, as microtubule separation may be caused by demembranation, it would be prudent to confirm these observations using more precise methods such as cryo-ET, which keeps tissues in a more accurate native state.

Among the most intriguing observations from our dataset is the “ciliary ring,” previously not described in any mammalian primary cilia. The low frequency of the ring suggest that it may not always survive de-membranization as in our procedure. Ring complexes around individual microtubules have been reported in budding yeast (*70*), namely the Dam1 complex that aids in microtubule stability under the control of Aurora kinase, but Dam1 rings are not seen in other fungal species nor are sequence homologues found in higher organisms. The ciliary ring that we describe is distinct from the ciliary necklace and bracelet, which have different morphology and localization, for example the ciliary bracelet has only been observed in the flagella of the unicellular green alga *Chlamydomonas* (*32*). Nor does it resemble the gamma-tubulin rings reported in *Xenopus*, whose function to cap the minus end and block MT growth (*71*). The only known ring-like structure that wraps around the entire axoneme is the membrane-bound septin ring at the cilium base (*72*). Septins are a family of GTP-binding proteins that act as diffusion barriers, and in primary cilia the septin ring is found below the transition zone. Thus location-wise, the septin ring location is too low to be the structure that we observe, which is microns above the cell surface. While the composition of the ring remains unknown, we suggest based on its morphology that this could be a tubulin-based complex that contributes to the structural integrity of the axonemal bundle. Having a mid-axonemal ring could support microtubule bundling, promote cilia assembly, and stabilize against disassembly. Whether the formation of the primary cilia ring, like the yeast Dam1 complex, is regulated by Aurora kinase signaling will be interesting to see, but beyond the scope of this study.

Finally, the observation of compound primary cilia raises questions about their purpose and the mechanism of numerical control. The majority of cells in human pancreatic islets contain a single primary cilium, consistent with the idea that there may exist evolutionary mechanisms to ensure single primary cilia number for proper function (*73*). While our images clearly demonstrate the existence of double and triple cilia, we do not know their functional significance, nor can we glean from surface scans their anatomical origin, for example whether they are joined together at the base or accompanied by extranumerary centrioles. The Polo-like kinase (PLK) family proteins are well-studied regulators for centriole biogenesis in human cancer cell lines, with PLK4 being an established driver of centrosome duplication (*74–76*). While these pathways have not been experimentally studied in pancreatic islets, a survey of published human single cell RNAseq datasets (*77, 78*) shows that all Plk1-5 genes including Plk4 have detectable and cell type-dependent expression in normal healthy human islets, suggesting that there indeed exist mechanisms for regulated cilia/centrosome duplication. The multi-cilia observed in our study were also healthy-appearing, with normal lengths and diameters and individual ciliary membranes, surrounded by a well-formed ciliary pocket. Thus we posit that, rather than being a diseased oddity, these multi-cilia may be a normal feature among islet cells, perhaps retained over evolution to maintain cilia density per unit surface area or volume. Given the heterogeneity of Plk gene expression, multi-ciliation may also be differentially regulated among islet cell subsets or be under hormonal control. Gain- and loss-of-function studies in the future might demonstrate a role for PLK or related factors in centrosome amplification, cilia biogenesis, and islet function.

## METHODS

### Islet preparation for SEM

Human islets were obtained from the Integrated Islet Distribution Program (IIDP). After retrieving islets from the shipping bottle, we washed and rested islets overnight in islet medium (RPMI 1640 with 11 mM glucose, 10% FBS, and 1% Penicillin-Streptomycin (Gibco, #15140122)). Islets were then plated on laminin-coated coverslips (treated with 0.5 μg laminin/cm^2^ for two hours), and allowed to adhere for three days. For fixation/extraction, islets were washed three times in warm PBS, then membrane-extracted with successive incubations, first for 5 minutes in cytoskeleton buffer (50 mM imidazole, 50 mM KCl, 0.5 mM MgCl2, 0.1 mM EDTA, 1 mM EGTA, pH 6.8) with 0.5% Triton X-100 and 0.25% glutaraldehyde, then for 10 minutes in cytoskeleton buffer with 2% Triton X-100 and 1% CHAPS. Islets were washed three times in cytoskeleton buffer before fixation overnight in a solution containing 2% glutaraldehyde in 0.15 M cacodylate buffer, pH 7.4 at 4 °C.

### Scanning electron microscopy

Coverslips were then transferred into 0.1% tannic acid in ultrapure water for 20 minutes at room temperature. Following this, samples were rinsed 4 times for 5 minutes each in ultrapure water and incubated in 0.2% aqueous uranyl acetate for 20 minutes. Samples were then rinsed 3 times for 10 minutes each in ultrapure water and dehydrated in a graded ethanol series (10%, 20%, 30%, 50%, 70%, 90%, 100% x3) for 5 minutes in each step. Once dehydrated, samples were loaded into a critical point drier (Leica EM CPD 300, Vienna, Austria) which was set to perform 12 CO_2_ exchanges at the slowest speed. Samples were then mounted on aluminum stubs with carbon adhesive tabs and coated with 5 nm of carbon and 5 nm of iridium (Leica ACE 600, Vienna, Austria). SEM images were acquired on a FIB-SEM platform (Helios 5 UX DualBeam Fisher Scientific, Brno, Czech Republic) using SEM imaging mode at 1.8 kV and 0.1 nA using TLD detector.

### Immunohistochemistry, confocal and super-resolution microscopy

Isolated human islets were washed with PBS and fixed with 4% paraformaldehyde (PFA) for 15 minutes and permeabilized with 0.3% Triton X-100 in PBS (PBST) for 10 minutes at room temperature. After incubation with blocking buffer (PBS with 10% normal goat serum) for 1 hour at room temperature, islets were incubated overnight at 4°C with primary antibodies diluted in PBST. Antibodies included mouse-anti-acetylated tubulin (Proteintech #66200-1-Ig, 1:400 dilution), rabbit-anti-ARL13b (Proteintech, 1:400 dilution), and mouse-anti-DNAIC1 (NeuroMab #75-372, 1:500 dilution). The next day, islets were washed, incubated with secondary antibodies for 1 hour at room temperature, and washed again in PBST. DAPI provided nuclear counterstain. Islets were mounted on glass slides with Prolong Gold Anti-fade (Thermo Fisher P36930) and imaged using an inverted Zeiss LSM880 and Ti2 Eclipse SIM microscope on Airyscan mode (Nikon, Ti-E).

### Morphometric analysis

Linear measurements of human islet cilia length and diameter were performed in ImageJ using the straight line or freehand line measurement tool. Intact full-length cilia images were selected for analysis, excluding those that were fragmented or disrupted during sample processing. Results were compiled in GraphPad Prism and calculated for average and standard deviation. Volume (V) and surface area (SA) calculations are derived by approximating the shape of the primary cilium to a truncated cone with base radius *r*_*1*_, tip radius *r*_*2*_, and height *h*, using the following equations:

For a circular truncated cone:

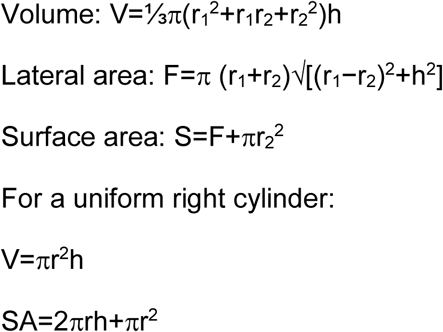

## ACKNOWLEDGMENTS

We are indebted to Dr. Ursula Goodenough along with Dr. Bob Schmidt, Robyn Roth, and the WashU Cilia Group for help with SEM data interpretation. We acknowledge the assistance of Drs. James Fitzpatrick and Praveen Krishnamoorthy at the Washington University Center for Cellular Imaging (WUCCI) in electron and light microscopy studies, supported by Washington University School of Medicine, The Children’s Discovery Institute of Washington University and St. Louis Children’s Hospital (CDI-CORE-2015-505 and CDI-CORE-2019-813), the Foundation for Barnes-Jewish Hospital (3770) and the Washington University Diabetes Research Center (DK020579). Human pancreatic islets were provided by the NIDDK-funded Integrated Islet Distribution Program (IIDP) (RRID:SCR_014387) at City of Hope, NIH grant #2UC4DK098085 and the JDRF-funded IIDP Islet Award Initiative. A.P. is supported by the Diabetes and Related Metabolic Diseases training grant T32DK007120. Other funding included NIH grants DK115795 and DK127748 to J.H. and DK115972 and DK123301 to D.P. Graphics were created with BioRender.com. The authors declare that they have no competing interests.

## Author Contributions

A.P., D.P., and J.H. conceptualized the study. A.P. and S.S. performed scanning electron microscopy. A.P. performed structured illumination microscopy. I.M. and J.H. performed morphometric analysis and immunohistochemistry. J.H.’s laboratory provided human islets. J.H wrote the manuscript with input from all authors.

## Notes

### Competing Interest Statement

The authors have declared no competing interest.

## REFERENCES

1. B. L. Munger, A light and electron microscopic study of cellular differentiation in the pancreatic islets of the mouse. Am J Anat. 103, 275–311 (1958).

2. L. Boquist, Cilia and vesicular particles in the endocrine pancreas of the Mongolian gerbil. J Cell Biol. 45, 532–541 (1970).

3. N. A. Zaghloul, N. Katsanis, Mechanistic insights into Bardet-Biedl syndrome, a model ciliopathy. J Clin Invest. 119, 428–437 (2009).

4. M. Fliegauf, T. Benzing, H. Omran, When cilia go bad: cilia defects and ciliopathies. Nat Rev Mol Cell Biol. 8, 880–893 (2007).

5. J. L. Badano, N. Mitsuma, P. L. Beales, N. Katsanis, The ciliopathies: an emerging class of human genetic disorders. Annu Rev Genomics Hum Genet. 7, 125–148 (2006).

6. S. Lodh, E. A. O’Hare, N. A. Zaghloul, Primary cilia in pancreatic development and disease. Birth Defects Research Part C: Embryo Today: Reviews. 102, 139–158 (2014).

7. P. diIorio, A. R. Rittenhouse, R. Bortell, A. Jurczyk, Role of cilia in normal pancreas function and in diseased states. Birth Defects Research Part C: Embryo Today: Reviews. 102, 126–138 (2014).

8. M. Titlbach, K. Fält, S. Falkmer, Postnatal maturation of the islets of Langerhans in sheep. Light microscopic, immunohistochemical, morphometric, and ultrastructural investigations with particular reference to the transient appearance of argyrophil insulin immunoreactive cells. Diabetes Res. 2, 5–15 (1985).

9. M. Yamamoto, K. Kataoka, Electron microscopic observation of the primary cilium in the pancreatic islets. Arch Histol Jpn. 49, 449–457 (1986).

10. N. Ashizawa, N. Hamamoto, T. Kaji, M. Watanabe, Scanning electron microscope examination of pancreatic ducts in stroke-prone spontaneously hypertensive rats (SHRSP). Int J Pancreatol. 22, 51–57 (1997).

11. A. A. Aughsteen, The ultrastructure of primary cilia in the endocrine and excretory duct cells of the pancreas of mice and rats. Eur J Morphol. 39, 277–283 (2001).

12. T. Iwanaga, T. Miki, H. Takahashi-Iwanaga, Restricted expression of somatostatin receptor 3 to primary cilia in the pancreatic islets and adenohypophysis of mice. Biomedical Research. 32, 73–81 (2011).

13. M. H. Greider, D. W. Elliott, Electron microscopy of human pancreatic tumors of islet cell origin. Am J Pathol. 44, 663–678 (1964).

14. E. A. Phelps, C. Cianciaruso, J. Santo-Domingo, M. Pasquier, G. Galliverti, L. Piemonti, E. Berishvili, O. Burri, A. Wiederkehr, J. A. Hubbell, S. Baekkeskov, Advances in pancreatic islet monolayer culture on glass surfaces enable super-resolution microscopy and insights into beta cell ciliogenesis and proliferation. Sci Rep. 7, 45961 (2017).

15. W. J. Gan, M. Zavortink, C. Ludick, R. Templin, R. Webb, R. Webb, W. Ma, P. Poronnik, R. G. Parton, H. Y. Gaisano, A. M. Shewan, P. Thorn, Cell polarity defines three distinct domains in pancreatic β-cells. J Cell Sci. 130, 143–151 (2017).

16. J. H. Cho, Z. A. Li, L. Zhu, B. D. Muegge, H. F. Roseman, E. Y. Lee, T. Utterback, L. G. Woodhams, P. V. Bayly, J. W. Hughes, Islet primary cilia motility controls insulin secretion. Science Advances. 8, eabq8486 (2022).

17. Z. A. Li, J. H. Cho, L. G. Woodhams, J. W. Hughes, Fluorescence imaging of beta cell primary cilia. Frontiers in Endocrinology. 13 (2022) (available at https://www.frontiersin.org/articles/10.3389/fendo.2022.1004136).

18. T. Caspary, C. E. Larkins, K. V. Anderson, The Graded Response to Sonic Hedgehog Depends on Cilia Architecture. Developmental Cell. 12, 767–778 (2007).

19. P. Kiesel, G. Alvarez Viar, N. Tsoy, R. Maraspini, P. Gorilak, V. Varga, A. Honigmann, G. Pigino, The molecular structure of mammalian primary cilia revealed by cryo-electron tomography. Nature Structural & Molecular Biology. 27, 1115–1124 (2020).

20. C. S. Xu, S. Pang, G. Shtengel, A. Müller, A. T. Ritter, H. K. Hoffman, S. Takemura, Z. Lu, H. A. Pasolli, N. Iyer, J. Chung, D. Bennett, A. V. Weigel, M. Freeman, S. B. van Engelenburg, T. C. Walther, R. V. Farese, J. Lippincott-Schwartz, I. Mellman, M. Solimena, H. F. Hess, An open-access volume electron microscopy atlas of whole cells and tissues. Nature. 599, 147–151 (2021).

21. S. Sun, R. L. Fisher, S. S. Bowser, B. T. Pentecost, H. Sui, Three-dimensional architecture of epithelial primary cilia. PNAS. 116, 9370–9379 (2019).

22. A. Müller, D. Schmidt, C. S. Xu, S. Pang, J. V. D’Costa, S. Kretschmar, C. Münster, T. Kurth, F. Jug, M. Weigert, H. F. Hess, M. Solimena, 3D FIB-SEM reconstruction of microtubule–organelle interaction in whole primary mouse β cells. J Cell Biol. 220, e202010039 (2020).

23. P. de Boer, N. M. Pirozzi, A. H. G. Wolters, J. Kuipers, I. Kusmartseva, M. A. Atkinson, M. Campbell-Thompson, B. N. G. Giepmans, Large-scale electron microscopy database for human type 1 diabetes. Nat Commun. 11, 2475 (2020).

24. J. W. Hughes, J. H. Cho, H. E. Conway, M. R. DiGruccio, X. W. Ng, H. F. Roseman, D. Abreu, F. Urano, D. W. Piston, Primary cilia control glucose homeostasis via islet paracrine interactions. PNAS (2020), doi:10.1073/pnas.2001936117.

25. C.-T. Wu, K. I. Hilgendorf, R. J. Bevacqua, Y. Hang, J. Demeter, S. K. Kim, P. K. Jackson, Discovery of ciliary G protein-coupled receptors regulating pancreatic islet insulin and glucagon secretion. Genes Dev. 35, 1243–1255 (2021).

26. M. V. Nachury, How do cilia organize signalling cascades? Philos Trans R Soc Lond B Biol Sci. 369, 20130465 (2014).

27. A. Fujioka, K. Terai, R. E. Itoh, K. Aoki, T. Nakamura, S. Kuroda, E. Nishida, M. Matsuda, Dynamics of the Ras/ERK MAPK cascade as monitored by fluorescent probes. J Biol Chem. 281, 8917–8926 (2006).

28. A. Benmerah, The ciliary pocket. Curr Opin Cell Biol. 25, 78–84 (2013).

29. R. Ghossoub, A. Molla-Herman, P. Bastin, A. Benmerah, The ciliary pocket: a once-forgotten membrane domain at the base of cilia. Biol Cell. 103, 131–144 (2011).

30. S. Sorokin, Centrioles and the formation of rudimentary cilia by fibroblasts and smooth muscle cells. J Cell Biol. 15, 363–377 (1962).

31. B. G. Barnes, Ciliated secretory cells in the pars distalis of the mouse hypophysis. J Ultrastruct Res. 5, 453–467 (1961).

32. N. B. Gilula, P. Satir, The ciliary necklace. A ciliary membrane specialization. J Cell Biol. 53, 494–509 (1972).

33. E. Tani, K. Ikeda, M. Nishiura, N. Higashi, Specialized intercellular junctions and ciliary necklace in rat brain. Cell Tissue Res. 151, 57–68 (1974).

34. C. Torikata, The ciliary necklace—A transmission electron microscopic study using tannic acid-containing fixation. Journal of Ultrastructure and Molecular Structure Research. 101, 210–214 (1988).

35. J. F. Reiter, O. E. Blacque, M. R. Leroux, The base of the cilium: roles for transition fibres and the transition zone in ciliary formation, maintenance and compartmentalization. EMBO reports. 13, 608–618 (2012).

36. Q. Hu, W. J. Nelson, Ciliary diffusion barrier: The gatekeeper for the primary cilium compartment. Cytoskeleton. 68, 313–324 (2011).

37. W. Breipohl, A. S. Mendoza, F. Miragall, Freeze-etching studies on the ciliary necklace in the rat and chick. J Anat. 130, 801–807 (1980).

38. S. Lee, H. Y. Tan, I. I. Geneva, A. Kruglov, P. D. Calvert, Actin filaments partition primary cilia membranes into distinct fluid corrals. Journal of Cell Biology. 217, 2831–2849 (2018).

39. C. Battle, C. M. Ott, D. T. Burnette, J. Lippincott-Schwartz, C. F. Schmidt, Intracellular and extracellular forces drive primary cilia movement. PNAS. 112, 1410–1415 (2015).

40. P. Kohli, M. Höhne, C. Jüngst, S. Bertsch, L. K. Ebert, A. C. Schauss, T. Benzing, M. M. Rinschen, B. Schermer, The ciliary membrane-associated proteome reveals actin-binding proteins as key components of cilia. EMBO reports. 18, 1521–1535 (2017).

41. H. J. Hoops, G. B. Witman, Outer doublet heterogeneity reveals structural polarity related to beat direction in Chlamydomonas flagella. J Cell Biol. 97, 902–908 (1983).

42. H. van den Hoek, N. Klena, M. A. Jordan, G. Alvarez Viar, R. D. Righetto, M. Schaffer, P. S. Erdmann, W. Wan, S. Geimer, J. M. Plitzko, W. Baumeister, G. Pigino, V. Hamel, P. Guichard, B. D. Engel, In situ architecture of the ciliary base reveals the stepwise assembly of intraflagellar transport trains. Science. 377, 543–548 (2022).

43. C. L. Williams, C. Li, K. Kida, P. N. Inglis, S. Mohan, L. Semenec, N. J. Bialas, R. M. Stupay, N. Chen, O. E. Blacque, B. K. Yoder, M. R. Leroux, MKS and NPHP modules cooperate to establish basal body/transition zone membrane associations and ciliary gate function during ciliogenesis. Journal of Cell Biology. 192, 1023–1041 (2011).

44. S. K. Dutcher, E. T. O’Toole, The basal bodies of Chlamydomonas reinhardtii. Cilia. 5, 18 (2016).

45. R. L. Weiss, D. A. Goodenough, U. W. Goodenough, Membrane particle arrays associated with the basal body and with contractile vacuole secretion in Chlamydomonas. Journal of Cell Biology. 72, 133–143 (1977).

46. P. M. Andrews, K. R. Porter, A scanning electron microscopic study of the nephron. Am J Anat. 140, 81–115 (1974).

47. A. S. Shah, S. L. Farmen, T. O. Moninger, T. R. Businga, M. P. Andrews, K. Bugge, C. C. Searby, D. Nishimura, K. A. Brogden, J. N. Kline, V. C. Sheffield, M. J. Welsh, Loss of Bardet–Biedl syndrome proteins alters the morphology and function of motile cilia in airway epithelia. Proc Natl Acad Sci U S A. 105, 3380–3385 (2008).

48. Y. Hou, G. J. Pazour, G. B. Witman, A Dynein Light Intermediate Chain, D1bLIC, Is Required for Retrograde Intraflagellar Transport. Mol Biol Cell. 15, 4382–4394 (2004).

49. D. Huangfu, K. V. Anderson, Cilia and Hedgehog responsiveness in the mouse. Proc Natl Acad Sci U S A. 102, 11325–11330 (2005).

50. L. Lester, C. Venditti, The prevalence of pili multigemini. Br J Dermatol. 156, 1362–1363 (2007).

51. M. Brissova, M. J. Fowler, W. E. Nicholson, A. Chu, B. Hirshberg, D. M. Harlan, A. C. Powers, Assessment of human pancreatic islet architecture and composition by laser scanning confocal microscopy. J Histochem Cytochem. 53, 1087–1097 (2005).

52. P. de Boer, B. N. Giepmans, State-of-the-art microscopy to understand islets of Langerhans: what to expect next? Immunology & Cell Biology. 99, 509–520 (2021).

53. E. Geron, S. Boura-Halfon, E. D. Schejter, B.-Z. Shilo, The Edges of Pancreatic Islet β Cells Constitute Adhesive and Signaling Microdomains. Cell Reports. 10, 317–325 (2015).

54. L. Orci, B. Thorens, M. Ravazzola, H. F. Lodish, Localization of the Pancreatic Beta Cell Glucose Transporter to Specific Plasma Membrane Domains. Science. 245, 295–297 (1989).

55. R. A. e Drigo, S. Jacob, C. F. García-Prieto, X. Zheng, M. Fukuda, H. T. T. Nhu, O. Stelmashenko, F. L. M. Peçanha, R. Rodriguez-Diaz, E. Bushong, T. Deerinck, S. Phan, Y. Ali, I. Leibiger, M. Chua, T. Boudier, S.-H. Song, M. Graf, G. J. Augustine, M. H. Ellisman, P.-O. Berggren, Structural basis for delta cell paracrine regulation in pancreatic islets. Nat Commun. 10, 1–12 (2019).

56. A. Molla-Herman, R. Ghossoub, T. Blisnick, A. Meunier, C. Serres, F. Silbermann, C. Emmerson, K. Romeo, P. Bourdoncle, A. Schmitt, S. Saunier, N. Spassky, P. Bastin, A. Benmerah, The ciliary pocket: an endocytic membrane domain at the base of primary and motile cilia. J Cell Sci. 123, 1785–1795 (2010).

57. J. B. Rattner, P. Sciore, Y. Ou, F. A. van der Hoorn, I. K. Y. Lo, Primary cilia in fibroblast-like type B synoviocytes lie within a cilium pit: a site of endocytosis. Histol Histopathol. 25, 865–875 (2010).

58. B. Mitchell, R. Jacobs, J. Li, S. Chien, C. Kintner, A positive feedback mechanism governs the polarity and motion of motile cilia. Nature. 447, 97–101 (2007).

59. I. Antoniades, P. Stylianou, P. A. Skourides, Making the connection: ciliary adhesion complexes anchor basal bodies to the actin cytoskeleton. Dev Cell. 28, 70–80 (2014).

60. M. Lemullois, C. Klotz, D. Sandoz, Immunocytochemical localization of myosin during ciliogenesis of quail oviduct. Eur J Cell Biol. 43, 429–437 (1987).

61. E. Boisvieux-Ulrich, M.-C. Lainé, D. Sandoz, Cytochalasin D inhibits basal body migration and ciliary elongation in quail oviduct epithelium. Cell Tissue Res. 259, 443–454 (1990).

62. W. F. Marshall, “Chapter 1 Basal Bodies: Platforms for Building Cilia” in Current Topics in Developmental Biology (Academic Press, 2008; https://www.sciencedirect.com/science/article/pii/S0070215308008016), vol. 85 of Ciliary Function in Mammalian Development, pp. 1–22.

63. J. Kim, J. E. Lee, S. Heynen, E. Suyama, K. Ono, K. Lee, T. Ideker, P. Aza-Blac, J. G. Gleeson, Functional genomic screen for modulators of ciliogenesis and cilium length. Nature. 464, 1048–1051 (2010).

64. V. Raghavan, Y. Rbaibi, N. M. Pastor-Soler, M. D. Carattino, O. A. Weisz, Shear stress-dependent regulation of apical endocytosis in renal proximal tubule cells mediated by primary cilia. Proceedings of the National Academy of Sciences. 111, 8506–8511 (2014).

65. R. Uzbekov, C. Prigent, Clockwise or anticlockwise? Turning the centriole triplets in the right direction! FEBS Letters. 581, 1251–1254 (2007).

66. P. Gönczy, Towards a molecular architecture of centriole assembly. Nat Rev Mol Cell Biol. 13, 425–435 (2012).

67. P. Satir, Chirality of the cytoskeleton in the origins of cellular asymmetry. Philos Trans R Soc Lond B Biol Sci. 371, 20150408 (2016).

68. S. Thitamadee, K. Tuchihara, T. Hashimoto, Microtubule basis for left-handed helical growth in Arabidopsis. Nature. 417, 193–196 (2002).

69. R. Orbach, J. Howard, The dynamic and structural properties of axonemal tubulins support the high length stability of cilia. Nat Commun. 10, 1838 (2019).

70. S. Westermann, A. Avila-Sakar, H.-W. Wang, H. Niederstrasser, J. Wong, D. G. Drubin, E. Nogales, G. Barnes, Formation of a dynamic kinetochore-microtubule interface through assembly of the Dam1 ring complex. Mol Cell. 17, 277–290 (2005).

71. Y. Zheng, M. L. Wong, B. Alberts, T. Mitchison, Nucleation of microtubule assembly by a γ-tubulin-containing ring complex. Nature. 378, 578–583 (1995).

72. Q. Hu, L. Milenkovic, H. Jin, M. P. Scott, M. V. Nachury, E. T. Spiliotis, W. J. Nelson, A Septin Diffusion Barrier at the Base of the Primary Cilium Maintains Ciliary Membrane Protein Distribution. Science. 329, 436–439 (2010).

73. M. R. Mahjoub, The importance of a single primary cilium. Organogenesis. 9, 61–69 (2013).

74. R. Habedanck, Y.-D. Stierhof, C. J. Wilkinson, E. A. Nigg, The Polo kinase Plk4 functions in centriole duplication. Nat Cell Biol. 7, 1140–1146 (2005).

75. J. Kleylein-Sohn, J. Westendorf, M. Le Clech, R. Habedanck, Y.-D. Stierhof, E. A. Nigg, Plk4-induced centriole biogenesis in human cells. Dev Cell. 13, 190–202 (2007).

76. D. K. Breslow, A. J. Holland, Mechanism and regulation of centriole and cilium biogenesis. Annu Rev Biochem. 88, 691–724 (2019).

77. S. Shrestha, D. C. Saunders, J. T. Walker, J. Camunas-Soler, X.-Q. Dai, R. Haliyur, R. Aramandla, G. Poffenberger, N. Prasad, R. Bottino, R. Stein, J.-P. Cartailler, S. C. Parker, P. E. MacDonald, S. E. Levy, A. C. Powers, M. Brissova, Combinatorial transcription factor profiles predict mature and functional human islet α and β cells. JCI Insight. 6, e151621 (2021).

78. J. Li, J. Klughammer, M. Farlik, T. Penz, A. Spittler, C. Barbieux, E. Berishvili, C. Bock, S. Kubicek, Single-cell transcriptomes reveal characteristic features of human pancreatic islet cell types. EMBO Rep. 17, 178–187 (2016).

